# Voluntary Running Sustains the Correction of Inflammation-Related Gene Expression Conferred by AAV Gene Therapy in Mdx Mice

**DOI:** 10.1101/2025.01.17.633557

**Authors:** Claire Yuan, Shelby E. Hamm, David L. Mack, Jean-Baptiste Dupont, Robert W. Grange

## Abstract

Duchenne Muscular Dystrophy is a fatal disease characterized by persistent skeletal muscle degeneration, inflammation, and fibrosis. Gene therapy using an adeno-associated virus derived vector and a microdystrophin transgene is currently under investigation in patients but the impact of physical activity on long-term therapeutic outcome remains poorly understood. Recently, we reported 21-weeks of voluntary wheel running complemented the positive endurance and muscle function outcomes of gene therapy in mdx mice. In the present study, we performed a transcriptomic analysis of the gene expression changes associated with functional recovery in the diaphragm. RNA-Sequencing and bioinformatic analysis revealed 2881 dysregulated genes in untreated and unexercised mdx mice including inflammatory and fibrotic signaling pathways frequently affected in Duchenne Muscular Dystrophy patients. Among the dysregulated genes, 774 were rescued towards WT after adeno-associated virus microdystrophin injection. Importantly, 93% of the rescued genes were maintained by voluntary running, which indicates that physical exercise has no significant impact on the outcome of gene therapy in the mdx diaphragm. Our study provides vital information that could help guide DMD patient follow-up protocols after treatment with gene therapy.

## Introduction

Duchenne Muscular Dystrophy (DMD) is a fatal X-linked neuromuscular disorder that affects 1 in 5000 males,^1^ and occurs due to both genetic inheritance and spontaneous mutations.^2^ The genetic mutations occur in one or multiple sites on the DMD gene which yields a truncated non-functional dystrophin protein.^3^ In the absence of a functional dystrophin, the dystrophin-glycoprotein complex (DGC) is also absent.^4^ The deficiency of the DGC from the sarcolemma results in susceptibility to mechanical stress and to disrupted signaling pathways.^4,5^ DMD patients experience muscular weakness during early childhood and the progressive loss of both ambulatory and cardio-respiratory functions.

The asynchronous and uncoordinated process of repairing dystrophic skeletal muscles forms a deleterious inflammatory cycle that drives dysfunctional dystrophic muscle (Fig. 1).^6^ Pro- and anti-inflammatory responses both play crucial roles in tissue healing, as the tightly regulated process of muscle regeneration after acute injury requires careful coordination between various immune cells.^7,8^ In chronic degenerative diseases such as DMD, membrane fragility is thought to facilitate persistent contraction-induced membrane damage which release endogenous damage-associated molecular patterns (DAMPS) and creatine kinase. This process can be described as ‘multiple-hit’, i.e., repeated bouts of muscle injury that create different stages of injury repair.^8^ Perpetual activation and recruitment of immune cell from the membrane damage leads to both a persistent pro-inflammatory and a wound healing macrophage phenotype that co-exist in the same environment.^8^ Activated immune cells have the capacity to release additional proinflammatory cytokines, such as Tumor Necrosis Factor-alpha (TNF-α), Interleukin-6 (IL- 6), and Interleukin-1 beta (IL-1β) that can cause additional cell lysis (Fig. 1).

**Figure 1.**
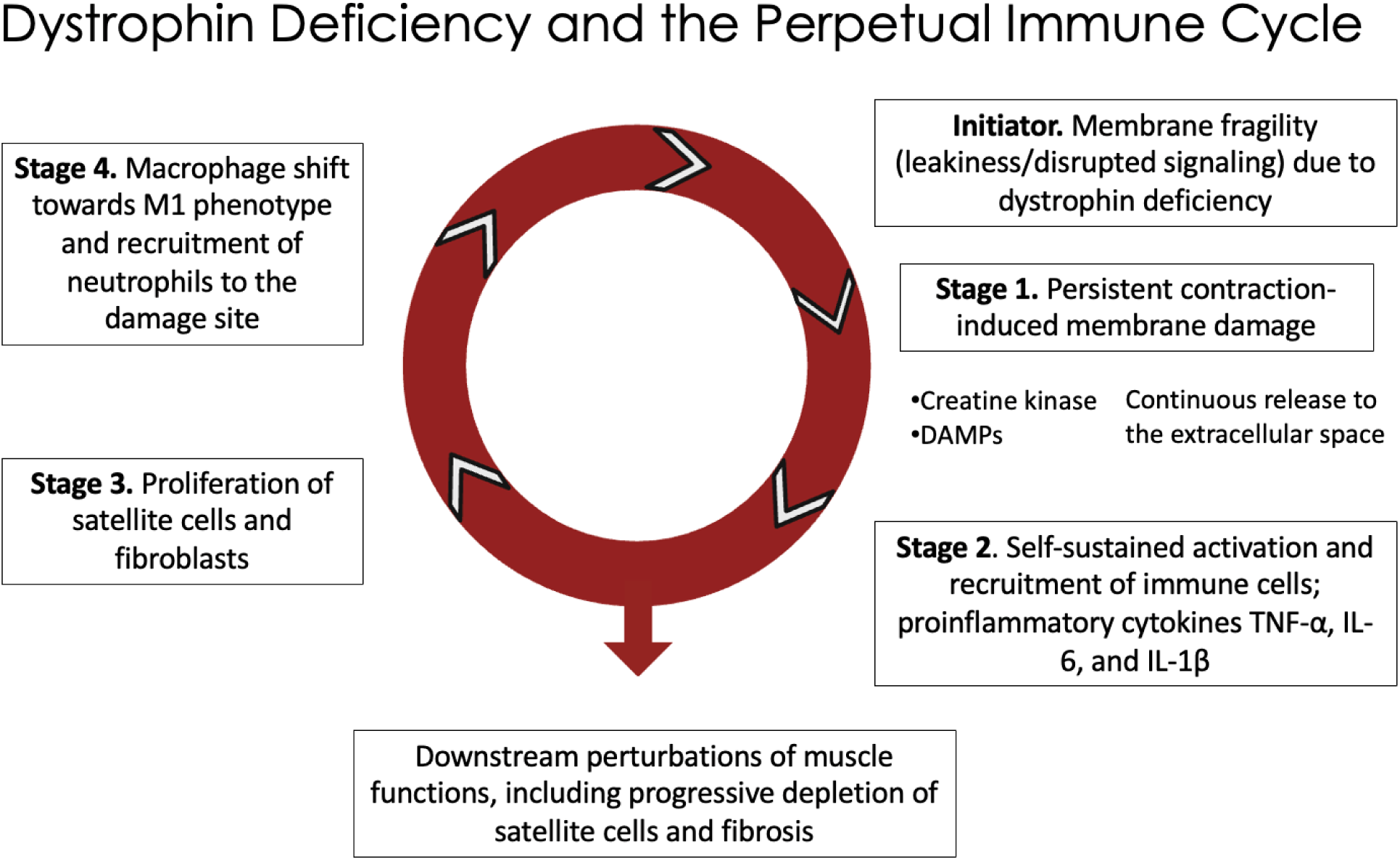
Dystrophin Deficiency and the Perpetual Immune Cycle. Dystrophic muscle membrane fragility induces immune cell infiltration and inflammatory responses.^8^ These responses exacerbate skeletal muscle necrosis, degradation, and fibrosis development resulting in a perpetual immune cycle.

As therapy strategies and treatments advance for DMD, functional improvement of diseased muscle is a potential outcome. Improvement of muscle function in individuals with DMD could lead to more opportunity for physical activity. Exercise, or prescribed physical activity, has been suggested as a potential therapeutic strategy to improve DMD muscle function and complement other treatments.^9–11^ However, muscle contraction increases membrane permeability in DMD skeletal muscle and exacerbates dystrophic pathophysiology.^12^ High-intensity exercise training such as treadmill running with a negative slope that elicits eccentric contractions induces acute tissue damage.^13^ In contrast, low-intensity voluntary wheel running for 21 weeks was reported to complement AAV microdystrophin gene therapy and enhance endurance capacity in mdx mice.^14^ In another report, voluntary wheel running was shown to trigger positive muscle adaptations such as increased muscle mass and a fiber type switch towards a slow contracting fatigue resistance phenotype.^15^ Furthermore, improved ability to run implies improved diaphragm function. While microdystrophin gene therapy has shown to improve endurance capabilities in dystrophic mice, the molecular signaling mechanisms behind this effect and the role of running to further enhance endurance are still undetermined. Although amelioration of inflammation is considered a potential treatment strategy for muscle wasting diseases, more study is needed to fully understand how inflammation contributes in chronic tissue degeneration and regeneration in a dystrophic phenotype.^16^ Furthermore, the rescue of inflammatory pathways by microdystrophin-based gene therapy and the additional impact of physical exercise needs to be carefully investigated.

In this study we focused on the inflammatory response in diaphragm and the impact of both adeno- associated virus (AAV) microdystrophin gene therapy (udys-GT) and voluntary exercise on this specific feature. The diaphragm was selected because it is considered the most severely affected muscle in mdx mice.^17,18^ Herein, we report the outcome of transcriptome analysis on the diaphragms obtained from our previous study.^14^

## Results

The groups described in Results are those defined in Hamm et al. 2021. Briefly, mdx mice were assigned to 3 groups: mdxRGT (run, gene therapy), mdxGT (no run, gene therapy), or mdx (no run, no gene therapy). Wild-type (WT) mice were assigned to WTR (run) and WT (no run) groups. Statistically differentially expressed genes were determined for each pair of groups as shown in Figure 2 : mdx vs WT, mdxGT vs WT, mdxRGT vs WT, mdxRGT vs WTR, mdxGT vs mdx, mdxRGT vs mdx, and mdxRGT vs mdxGT.

**Figure 2.**
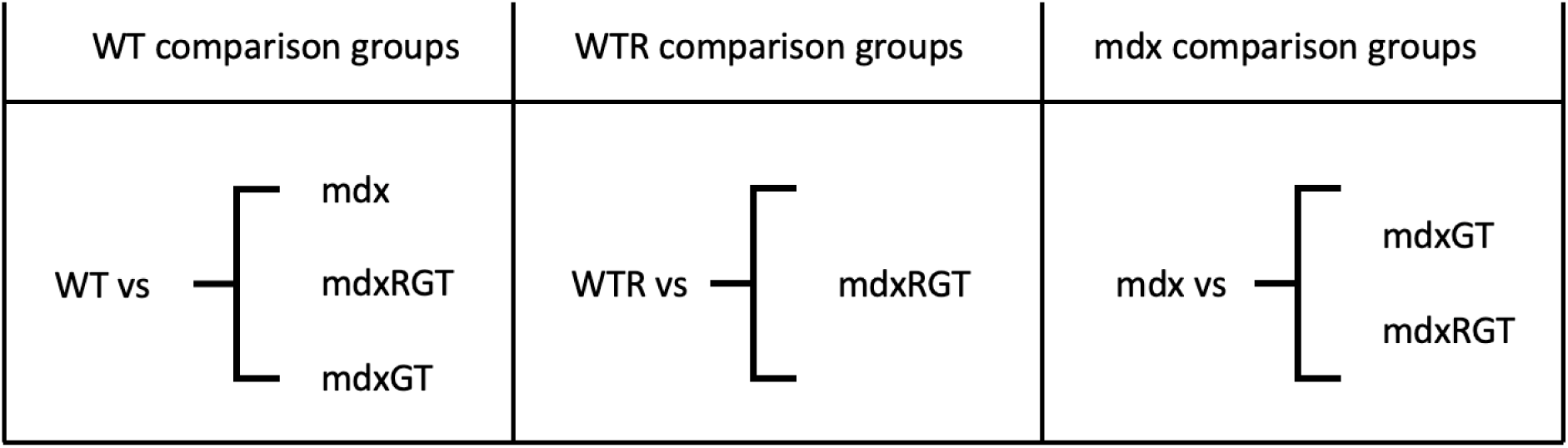
Schematic showing the treatment groups in pairs used for the differentially expressed gene analysis. The different comparison groups were further analyzed for statistical significance of gene expression.

We first analyzed the differentially expressed genes (DEGs) in the untreated unexercised mdx group, in comparison to the WT control group. We found 2881 DEGs with a |log2 (fold-change)| > 1 & P-adj < 0.05 (Fig. 3a). Gene ontology (GO) analysis of the biological processes significantly enriched in this DEG list pointed to metabolic processes (“fatty-acid beta-oxidation using acyl-CoA”, “mitochondrial electron transport”, “oxidative phosphorylation”), the inflammatory response (“interleukin-1 production”, “neutrophil activation”) and skeletal muscle biology (“skeletal muscle contraction”, “muscle system process”), confirming known phenotypes associated with the Duchenne pathology in mdx mice (Fig. 3b). Next, a Principal Component Analysis (PCA) was applied to all samples by considering the 1000 most variable genes in the dataset, and it showed clear separation of the 5 groups along Dimension 1, accounting for 57 % of the variance (Fig. 3c). WT and WTR groups are observed to be clustered together, indicating that voluntary exercise has a minor impact on the diaphragm in WT mice compared with the perturbations induced by the disease or its rescue. The mdxGT group appears closer to WT and WTR than to mdx, suggesting a beneficial effect of the gene therapy treatment. The mdxRGT group is clustering close to mdxGT, but the resolution of the PCA does not allow precise characterization of the effects of running on the transcriptome after gene therapy.

**Figure 3.**
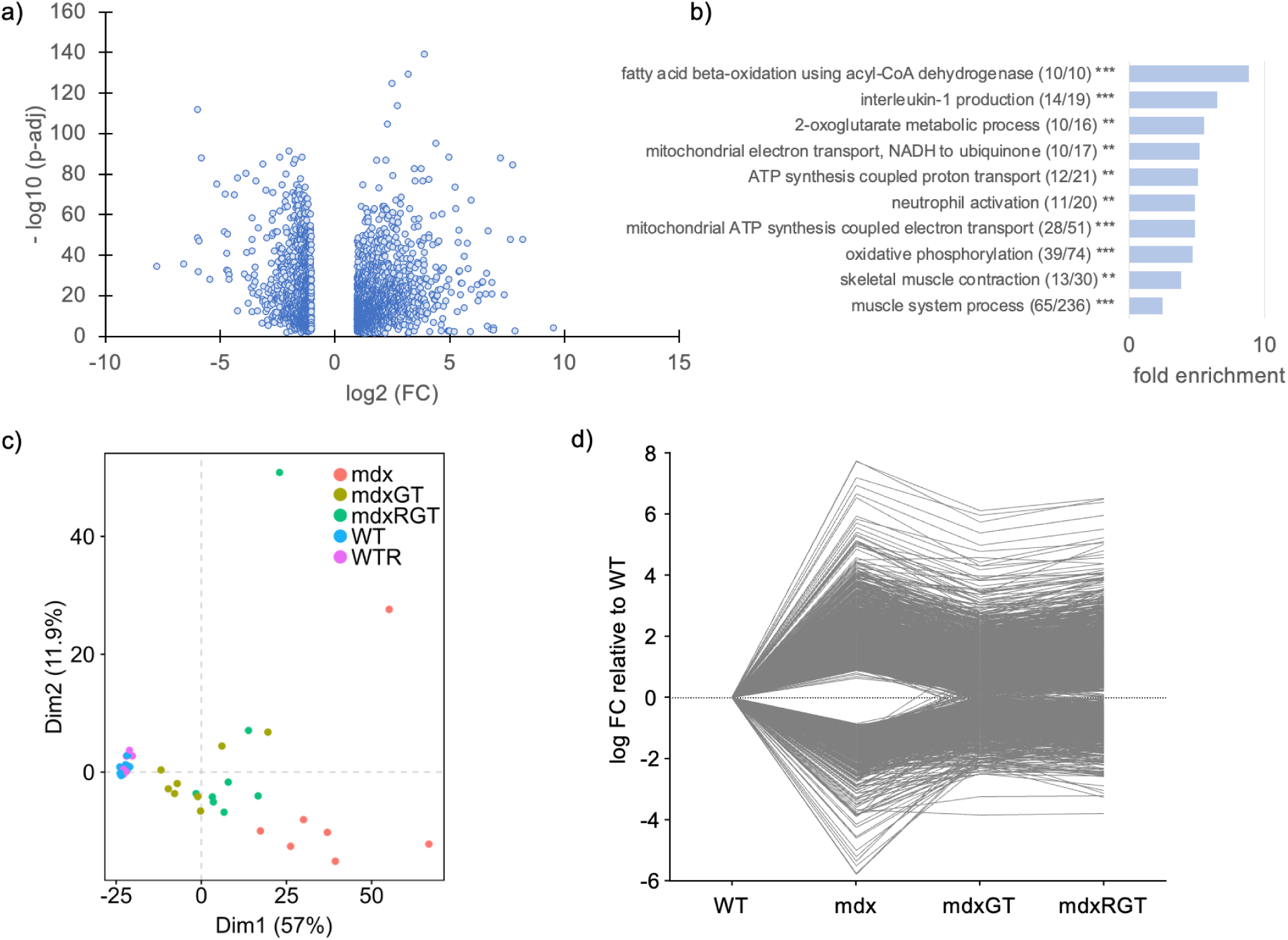
An overview of the RNA sequencing results. (a) Volcano plot of differentially expressed genes in mdx mice vs. WT controls (|log2 (fold-change)| > 1 & P-adj < 0.05). (b) Gene Ontology – Biological Process on differentially expressed genes in mdx vs WT. Gene ontology analysis showing the biological process enriched using all the DEGs between mdx and WT genes, with a particular focus on pathways involved in inflammation, such as IL-1 related pathways. Ratios between brackets indicate the number of genes found both in the ontology and in the list of DEGs, relative to the total number of genes listed in the ontology **p<0.01; ***p<0.001; (c)Principal Components Analysis of the RNA Seq data in the diaphragm showing the variation of the top 1000 genes between samples. Each dot represented a mouse with color differentiated by their treatment group. Dim1 represent principal component 1 and Dim2 represent principal component 2. (d) The relative expression of the genes dysregulated in the diaphragm of mdx compared to WT, and the effect of microdystrophin gene therapy +/-run. The gene expressions are plotted in log fold change (log FC).

Finally, we examined the overall transcriptome remodeling in the three groups of mdx mice (mdx, mdxGT and mdxRGT) by plotting the log fold-change (FC) values of the DEGs relative to the WT group (Fig. 3d). In agreement with the PCA, we observed that gene therapy globally rescues the log FC values of both upregulated and downregulated DEGs towards 0, indicating that the expression of these genes gets closer to the WT group after AAV injection. Although the level of rescue is very close in the mdxGT and mdxRGT groups, it seems slightly higher without voluntary exercise.

To identify genes that were rescued by microdystrophin and remained rescued if running was combined with microdystrophin, the following custom filters were used (Fig. 4). Filter 1: The first filter discriminated dysregulated genes due to the mdx phenotype when compared to WT (fold-change > 1, p-value < 0.05, N = 2881 genes). Filter 2: The second filter discriminated the genes identified by filter 1 which are also differentially expressed in the mdxGT vs mdx comparison and demonstrated rescued gene expression towards the WT (total number of genes = 774). There were 289 genes among the 774 dysregulated genes rescued by gene therapy that were downregulated in mdx compared to WT. After gene therapy was applied in mdxGT, they were all upregulated compared to mdx. There were 485 genes upregulated in mdx compared to WT. All were downregulated after gene therapy was applied. Filter 3: The third filter identified the rescued genes that met the previous two criteria (Filters 1 and 2) and that were not differentially expressed in the mdxRGT vs mdxGT comparison group (indicated as N/A; total number of genes = 723). After application of these 3 filters, 723 genes in the mdxGT group were unchanged by running in the mdxRGT group. This outcome suggests that at the transcriptome level, running preserved the effects of gene therapy based on the lack of gene expression differences in the mdxRGT vs mdxGT among the majority (93.41%, 723 out of 774 genes) of the genes rescued by gene therapy. Only 6.59% (774 – 723 = 51) of the rescued genes were altered by running. Among the rescued genes preserved by running, 18.26% (132 out of 723 genes) were completely rescued towards the WT or WTR level in both mdxGT and mdxRGT. Completely rescued genes were categorized by a lack of gene expression differences in mdxGT vs WT and mdxRGT vs WTR. Altogether, this analysis shows that running has little impact on the extent of transcriptome rescue by AAV microdystrophin gene therapy in the mdx diaphragm.

**Figure 4.**
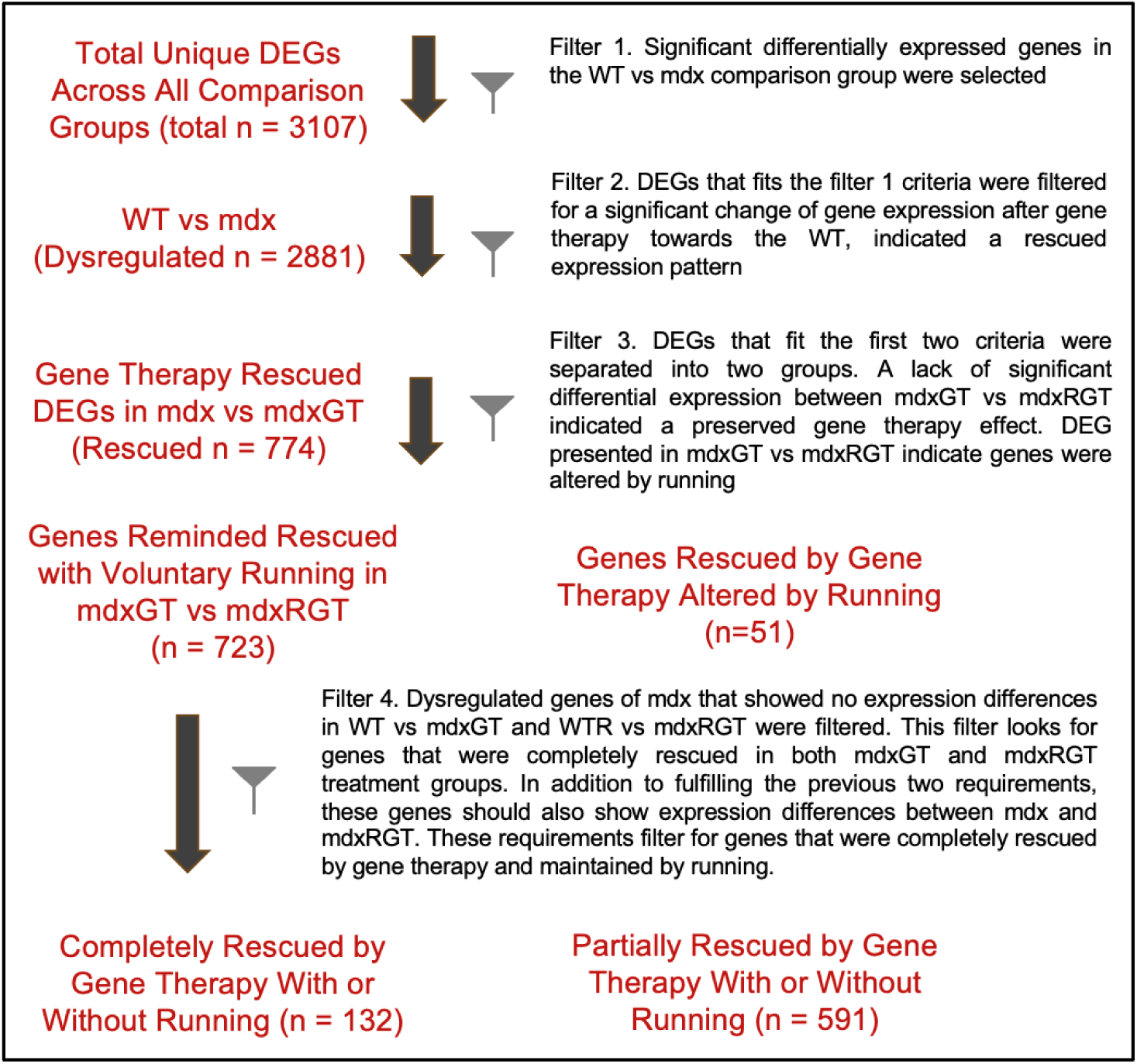
Workflow of the DEG analysis. The RNA Seq results of n = 3107 unique DEGs were categorized using R into these subgroups: Dysregulated, n = 2881: The dysregulated genes in mdx (significant differential gene expression in mdx compared to WT). Among the 2881 dysregulated genes, gene therapy rescued 774 of them towards the WT (significant differential gene expression in mdxGT compared to mdx and in the opposite direction of gene expression from mdx vs WT comparison group). When voluntary wheel running was applied in concert to gene therapy, 723 out of the 774 rescued genes maintained the rescued state (not significantly different in mdxRGT vs mdxGT group). While 51 genes rescued by gene therapy was altered by running in mdxRGT (gene expression in mdxRGT significantly different from mdxGT). The 774 genes were further categorized into another 2 subgroups based on their presence in the mdxGT vs WT, mdxRGT vs WTR and mdxRGT vs mdx group. Lack of DEGs in mdxGT vs WT and mdxRGT vs WTR but has significant expression differences in mdxRGT vs mdx group were considered as rescued (n=132). Otherwise, they were categorized as partially rescued (n=591).

To further examine the extent of the rescued effect, the WT and WTR groups were compared with mdxGT and mdxRGT mice, respectively, using the expression values of the 723 genes rescued by gene therapy and not affected by running. Genes that showed no significant expression difference in both mdxGT vs WT and mdxRGT vs WTR were categorized as completely rescued. It should be noted that genes that did not show expression differences when comparing mdxRGT to mdx were excluded during this analysis. Among the 723 genes, 132 genes were determined to be completely rescued. The other 591 genes that did not meet these requirements were categorized as partially rescued.

Analysis of dysregulated signaling pathways in mdx vs WT comparison group To further characterize the pathways involved in the mdx diaphragm and rescued by gene therapy with and without running, we used the Ingenuity Pathway Analysis (IPA) suite. IPA analyses on the genes dysregulated in the mdx diaphragm showed that inflammation is a key contributor to the DMD pathology at this time point (Fig 4a).

From the 2881 DEGs between mdx and WT mice, the IPA Comparison Analysis predicted 133 dysregulated canonical pathways from DEGs in mdx compared to WT. Of these 133, 117 pathways were predicted to be activated and 16 were predicted to be inhibited. Subsequently, we investigated the modulation of signaling pathways using the 723 dysregulated genes rescued by gene therapy and maintained by running. We found 39 canonical pathways predicted to be activated in the mdx vs WT group based on IPA and 1 predicted to be inhibited (Fig. 5a). Pathways were presented as hierarchical clustering (z > |2| & p <0.05 (−log 10 (P-value) > 1.3)). A total of 40 dysregulated signaling pathways met the criteria for significance in mdx vs WT, with 39 pathways activated and one pathway inhibited. Gene therapy rescued (inhibited) 38 of these pathways, thus affirming the therapeutic influence of gene therapy on signaling pathways towards WT activity. It is worth noting that some of these pathways remain activated in the mdxGT vs WT comparison group, implying that some of the genes involved in the signaling pathways were only partially rescued. Hierarchical clustering was used to visualize the similar expression pattern of the signaling pathways across different treatment groups. We further investigated the signaling pathways to determine the biological process they are involved in. Some of the notable activated pathways in mdx vs WT points towards immune response such as natural killer cell signaling, T cell receptor signaling and Toll-like receptor signaling (Fig 4c). To understand the role of the immune response with AAV microdystrophin gene therapy, we looked at the most activated pathway in mdx vs WT and also the most inhibited pathway in mdxGT vs mdx using IPA analysis. This pathway was the Pathogen Induced Cytokine Storm Signaling Pathway (PICSSP). Several inflammatory cytokines known for their function in immune cells recruitment were part of the signaling pathways, such as IL-6 and IL-1β.

**Figure 5.**
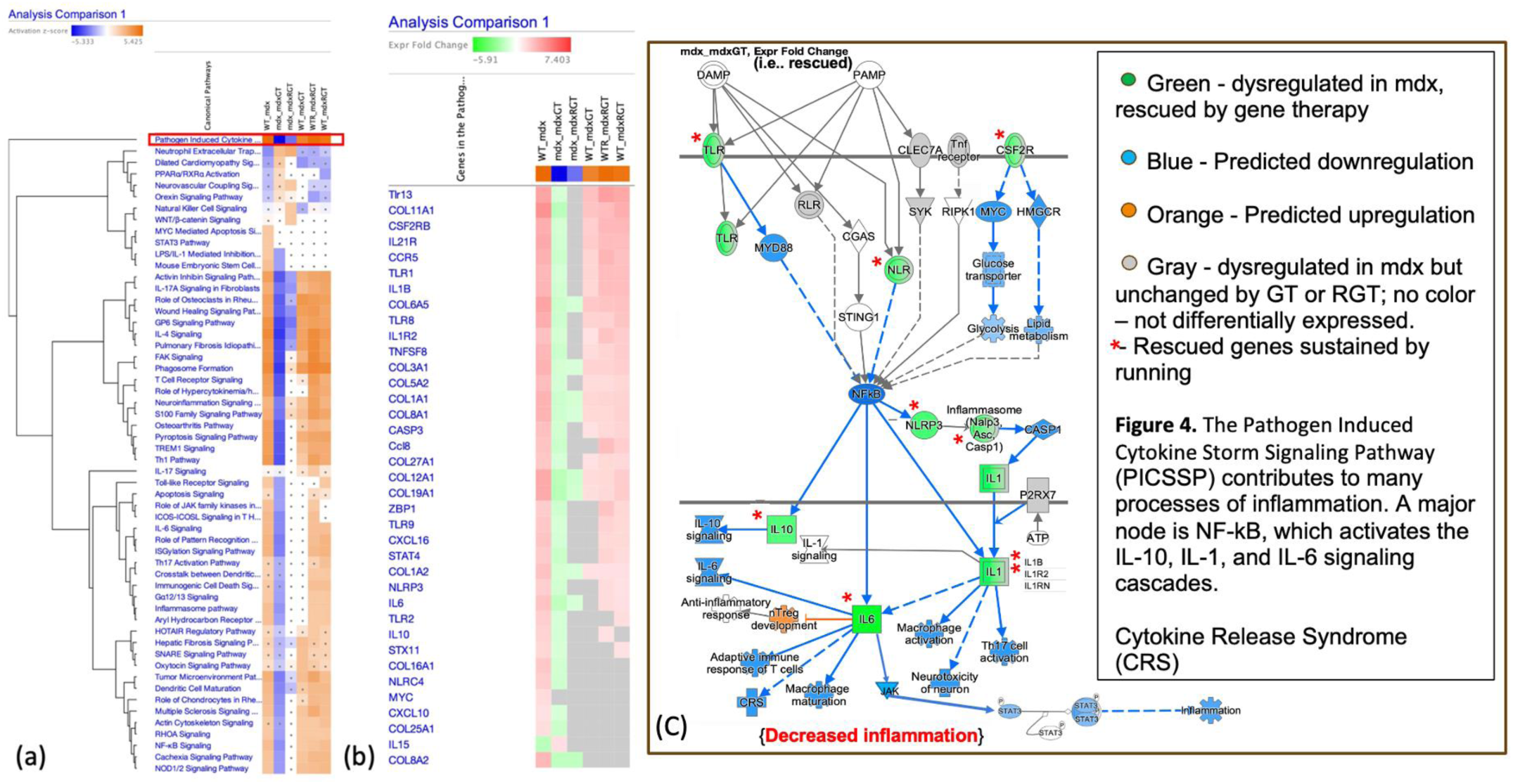
Signaling pathway enrichment detected using IPA. (a) A heat map of all the signaling pathways activated or inhibited across all comparison group using the genes that were rescued and maintained (n=723). The top signaling pathway activated in mdx compared to WT is the Pathogen Induced Cytokine Storm Signaling Pathway (PICSSP), outlined in red. (b) A list of rescued and maintained genes expressed in the activation of PICSSP in mdx. (c) PICSSP genes predicted to be downregulated by gene therapy and maintained by running. Genes that were dysregulated but not rescued by gene therapy were also included in this pathway and colored grey by IPA. We included these non-rescued genes in the pathway analysis to provide a comprehensive view of the effect of dystrophic signaling process and the effect of gene therapy. Lastly, red asterisks were placed next to the rescued genes preserved after running wheel activity.

Among the 37 genes involved in the PICSSP pathway, 36 were upregulated in mdx compared to WT and one – coding for interleukin 15 - was downregulated in mdx (Fig. 5(b)). The PICSSP pathway was then carefully analyzed in the light of the “rescued” status of its founder genes (Fig. 5 (c)). The inhibition of inflammatory signaling pathways revealed during IPA analysis highlighted the biological significance of gene therapy in mdx mice. Running did not impact the rescue of these genes, thereby maintaining the rescued signaling pathways.

Overall, our analysis has shown that signaling molecules of the inflammatory pathways were rescued towards the wild type in AAV microdystrophin gene therapy treated mdx mice. In addition, voluntary wheel running sustained the majority of rescued inflammatory gene expression in AAV microdystrophin treated mdx mice e.g., mdxRGT compared to mdxGT groups.

## Discussion

The impact of physical exercise in young DMD boys treated by AAV gene therapy is an important question to be answered, as various AAV vectors expressing microdystrophin transgenes are in advanced clinical stages. Our group recently reported the effects of voluntary wheel running on the physiological functions of microdystrophin AAV udys-GT treated mdx mice.^14^ The endurance capabilities assessed using treadmill running fatigue time in mdx were rescued towards the wild type after udys-GT treatment and maintained with voluntary wheel running.^14^ Skeletal muscle function assessed using in vivo plantarflexor torque showed an improvement in normalized torque in mdxRGT when compared to mdxGT.^14^ The quadriceps microdystrophin expression level of the udys-GT treated mice were increased with access to running wheel. However, microdystrophin expression level and power output in the diaphragm were decreased with running. The modulation of gene expression of udys-GT treated mdx mice with or without voluntary wheel running could shed light on the impact of exercise on the microdystrophin treatment. Herein, we investigated the overall changes in gene expression in response to gene therapy and exercise in the diaphragm of mdx mice and the associated signaling pathways. Overall, both the upregulated and downregulated genes were rescued by gene therapy towards the WT level in the mdx mice and running maintained the rescue profile for the majority of these genes.

After rigorous filtering of the DEGs and detailed analysis with IPA, we focused on the Pathogen- Induced Cytokine Storm Signaling Pathway (PICSS) because it includes inflammatory pathways known to be dysregulated in both DMD patients and mdx mice, primarily , the nuclear factor kappa-light-chain- enhancer of activated B cells (NF-kB) and the Interleukin-6/Janus Kinase/Signal Transducer and Activator of Transcription (IL-6/JAK/STAT) signaling pathways. The IL-6 activated signaling pathway is translationally relevant because IL-6 concentration is elevated in both DMD patients and at the necrotic stage of mdx mice.^19–22^ DMD patient IL-6 concentration level was reported higher compared to the healthy control.^21^ It is interesting to note that among the three inflammatory mediators (IL-6, IL-1β, and TNF-α), IL-6 transcripts were reported to be significantly downregulated in the diaphragm of 24 week-old mice compared to 4 week-old mdx mice.^22^ However, the expression of IL-6 at both age points were significantly higher compared to WT mice.^22^ Consistent with this finding, the diaphragm of 29- week-old mdx mice in our data set showed a significant upregulation compared to WT mice (log 2-fold change = 2.15). Gene expression of the TLR downstream signaling pathways, such as IL-1β, was significantly upregulated in exacerbated pathology of physically active mdx mice.^23^ These proinflammatory cytokines contribute to the recruitment and maturation of immune cells and perpetuate the detrimental inflammatory environment in dystrophic skeletal muscles. In our data, TLR-2, IL-6 and IL-1β were shown to be rescued by gene therapy and voluntary wheel running did not exert a detectable expression change.

Aberrant activation of transcription factors such as JAK/STAT and NF-kB were reported in the diaphragm of mdx mice at 6 weeks old.^24^ Specifically, STAT4 and myc protein were increased.^24^ However, the ablation of STAT3 in the satellite cells of mdx mice exacerbated fibrosis and inflammation indicate the complicated role STAT plays in cellular signaling and homeostasis.^25^ It is possible that acute pathway activation promotes muscle regeneration but elevated and persistent expression of IL-6 or complete depletion of STAT3 is detrimental to the regeneration process in the muscle microenvironment.

Pathological inflammation and fibrosis are induced by the release of DAMPS and creatine kinase due to membrane instability. ECM derived DAMPs such as tenascin C, and fibronectin are proteins located in the extracellular matrix. They can serve as signaling molecules to elicit inflammatory response through proteolysis by proteolytic enzymes. Tissue injury promotes the release of ECM DAMPs and some were reported to be dysregulated in mdx mice subjected to voluntary wheel running and in DMD patients.^23^ In our data, tenascin C, and fibronectin were also upregulated in mdx mice compared to the WT. Gene therapy rescued their expression towards the WT level which remained unchanged after wheel running.

Overall, our analysis suggests that physical activity conserved the gene expression profiles associated with positive microdystrophin gene therapy effects, without exacerbating the immune response and fibrosis formation in the mdxRGT mice. This outcome supports our previous hematoxylin and eosin histological observations that fibrotic tissue and immune cell infiltration are reduced in gene therapy treated mdx mice with or without running compared to mdx.^14^ A lack of significant differential expression of these genes mentioned above in mdxRGT mice could indicate that gene therapy rescued the muscle fibrosis phenotype in the mdx mice, and this therapeutic effect was preserved after 21 weeks of physical activity.

Exercise can be either beneficial or detrimental depending on the modality used, the intensity, duration, and frequency of the activity, disease state, and the age of the mdx mice.^15^ If regular physical activity can modulate these signaling pathways in a beneficial manner, then there might be value in adding physical activity as part of the treatment in concert with gene therapy. The signaling pathway discussed above should be monitored if physical activity is explored as part of a treatment strategy in the future.

## Materials and Methods

### Diaphragm tissue

Diaphragms were obtained from the mice used in the Hamm et al., 2021 study. All experiments performed on mice in the Hamm et al., 2021 study were approved by the Institutional Animal Care and Use Committee at Virginia Tech. Briefly, 20 WT (C57BL-10ScSnJ/Jax strain #000476) and 20 mdx mice (C57BL/10mcSn-DMDmdx/Jax strain #001801) were divided into 5 groups (each n=8) at 7 weeks of age. Mice from the gene therapy groups were injected with an AAV9 microdystrophin vector in the tail vein at 2e14 vg/kg and all other groups received an equivalent volume of excipient. Over the study duration of 21 weeks, WTR and mdxRGT groups were housed in cages with a functional running wheel. WT, mdx, and mdxGT were housed in cages with a locked running wheel. Thirty-eight mice completed the study, with one WT and one mdx mouse dying early from dehydration ^14^ At the conclusion of the study, all mice were deeply anesthetized with ketamine/xylazine and diaphragms excised and stored in RNALater at -80°C.

### RNA Seq Analysis

RNA seq library preparation and sequencing were performed as described (Hamm et al. 2021, Supplemental Methods). Raw counts were generated using Genewiz mapped to the Mus musculus GRCm38 reference genome. EdgeR operated on the open-source RStudio environment for R was used for the secondary analysis of the data, including principal component analysis (PCA), heatmap and differentially expressed genes (DEGs). The code used to generate normalized counts to compare between groups are listed in detail (Dupont et al., 2020, Supplemental Information). Gene expression with |log2 (fold-change)| > 1 & P-adj < 0.05 discriminated the differential expression of a given gene between groups.

Gene ontology results were produced using the Gene Ontology (GO) Enrichment Analysis from GO Consortium online platform (https://geneontology.org/). GO plot and volcano plot of mdx compared to WT were generated using MICROSOFT Excel. RNA transcripts were analyzed in the diaphragm for 5 different treatment groups: WT, WTR, mdx, mdxGT, mdxRGT. After DEGs were imported into R, filters were programmed to query genes of interest based on these criteria:

1. The “Dysregulated Genes” in mdx were indicated by DEGs present in the mdx vs WT group filtered by "!is.na” in R .
2. The “Gene Therapy Rescued Dysregulated Genes” in mdxGT were indicated by DEGs present in both the mdx vs WT and mdxGT vs mdx comparison group. Specifically, genes were shown to be upregulated (positive) in mdx vs WT should be downregulated (negative) in the mdxGT vs mdx group to indicate a rescued effect, and vice versa; downregulated genes in the mdx vs WT group shown to be positive in the mdxGT vs mdx group would be considered as rescued.
3. In addition to meeting the previous 3 criteria, “rescued genes maintained by running” (Maintained) were determined by the lack of DEGs (indicated by N/A) in the mdxRGT vs mdxGT group. Alternatively, existing DEGs in this category indicate the effect of gene therapy was altered by running.
4. The Maintained genes were further categorized into two subgroups: completely rescued and partially rescued. This was using the “is.na” in R applied in mdxGT vs WT and WTR vs mdxRGT group to look for genes lack of significant expression differences in these two groups. In addition, there is a significant difference in mdx vs mdxRGT group. Genes that fit these criteria were considered completely rescued, otherwise they are considered partially rescued.

The canonical molecular signaling pathways associated with these genes were identified by Qiagen IPA Ingenuity Pathway Analysis (IPA).

## Author Contribution

Conceptualization, R.W.G.; methodology, R.W.G.; investigation, R.W.G., S.E.H. and C.Y.; data curation, R.W.G., S.E.H., J.B.D., C.Y., and D.L.M.; formal analysis, R.W.G., J.B.D., S.E.H., C.Y., and D.L.M.; resources, R.W.G.; writing – original draft, R.W.G., and C.Y.; writing – review & editing, R.W.G., J.B.D., C.Y., and D.L.M.; visualization, R.W.G., J.B.D., C.Y., and D.L.M.; supervision, R.W.G. and J.B.D.; resource, R.W.G. and J.B.D.; project administration, R.W.G.; funding acquisition, R.W.G.

## Funding

Provided by Solid Biosciences Inc.

Data Availability Statement

The RNA-seq datasets will be deposited at the Gene Expression Omnibus (GEO) database and will be made publicly available as of the date of publication.

